# Automatic genome segmentation with HMM-ANN hybrid models

**DOI:** 10.1101/034579

**Authors:** Li Shen

**Affiliations:** Department of Neuroscience, Icahn School of Medicine at Mount Sinai, New York, NY 10029, USA

## Abstract

We consider the problem of automatic genome segmentation (AGS) that aims to assign discrete labels to all genomic regions based on multiple ChIP-seq samples. We propose to use a hybrid model that combines a hidden Markov model (HMM) with an artificial neural network (ANN) to overcome the weaknesses of a standard HMM. Our contributions are threefold: first, we benchmark two approaches to generate targets for ANN training on an example dataset; second, we investigate many different ANN models to identify the ones with best predictions on chromatin states; third, we test different hyper-parameters and discuss how they affect the machine learning algorithms’ performance. We find our best performing models to beat two pervious state-of-the-art methods for AGS by large margins.

## 1 Introduction

The human genome contains a huge collection of functional elements that play key roles in regulating health and diseases. Identifying these functional elements is like labelling the various genomic locations with different colors and each color corresponds to a distinct functional category. This has been a very slow and painstaking process for experimental biologists. Chromatin marks, such as transcription factors, histone modifications and DNaseI hyper-sensitive sites, have been associated with many important cellular processes and diseases. Researchers hypothesize that combinations of the chromatin marks, or so-called chromatin states, are linked to distinct biological functions^1^. Each chromatin state is associated with genomic locations that may correspond to specific types of functional elements, such as transcriptional start sites (TSSs), enhancers, transcribed and repressed regions. With the advent of the next-generation sequencing (NGS) technology and particularly ChIP-seq, the genomic locations of a multitude of chromatin marks have been mapped, generating a large amount of data^2,3^.

Each ChIP-seq sample can be used to derive the enrichment values of a chromatin mark on the whole genome at single-base resolution, which provides an opportunity to use machine learning techniques to exploit the rich set of ChIP-seq data that has already been generated in public. Automatic genome segmentation (AGS) aims to cluster multiple ChIP-seq samples based on their combinations and assign meaningful labels to them. One choice is to use the HMM that is widely used to model biological sequences. The HMM can be used to assign a discrete state number to a genomic location without being given any training labels, basically performing an unsupervised clustering for all genomic regions. This approach was previously explored^4^ and was later extended into a dynamic Bayesian network model that makes single-base resolution inference computationally feasible^5^. However, standard HMMs suffer from several weaknesses that can lead to poor discrimination. Here, we consider the use of a hybrid model that combines an ANN with an HMM to overcome its limitations. This “hybrid” approach had been explored to enhance automatic speech recognition (ASR) for many years^6^ but only recently became mainstream^7^. As far as we know, we are the first group to explore this approach for AGS.

We consider two approaches to hypothesize the state labels from ChIP-seq data as targets for ANN training and benchmark them on an example dataset. We test many different ANN models to identify the ones with the best predictions on chromatin states and find that a wide but shallow network may be most suitable for our inputs. We also discuss how the hyper-parameters used in our machine learning algorithms affect the system’s performance. Finally, we find our best performing models to beat two previous approaches by large margins.

## 2 Approach

### 2.1 Overview of the HMM-ANN hybrid model

The genomic ChIP-seq data can be represented as 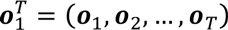 where 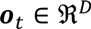 is a vector of signals at a fix-sized window (indexed by t) from D ChIP-seq samples. In this study, we use Gaussian mixture models (GMMs) as the probabilistic distributions for ***o***_*t*_. The window size is chosen to be 200bp across this study. The HMM specifies a generative model that the signals are generated from a sequence of hidden states 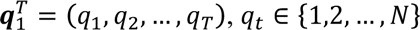 are discrete numbers representing chromatin states. The HMM is characterized by the following parameters: initial probabilities *π*_*i*_ = *p*(*q*_1_ = *i*); transition probabilities *a*_*ij*_ = *p*(*q*_*t*+1_ = *j*|*q*_*t*_ = *i*); observation probabilities *b*_*j*_(***o***_*t*_ = *p*(***o***_*t*_|*q*_*t*_ = *j*);. In a standard HMM, the ***o***_*t*_ is only dependent on the *q*_*t*_ but not any of the neighboring observations. This certainly contradicts with the biological reality. However, trying to directly model the relationships between ***o***_*t*_ and the neighboring observations and/or hidden states will involve an exponentially growing number of parameters. A hybrid model was initially proposed to circumvent this problem in ASR, which can be seen by applying the Bayes rule so that 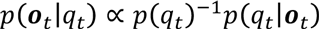 where the prior probabilities *p*(*q*_*t*_)can be empirically estimated and the posterior probabilities *p*(*q*_*t*_|***o***_*t*_) can be approximated by a classifier. The positeline/probabilities can often be better approximated by adding a contextual window: 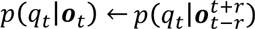 where *r* is the radius, which allows the neighboring information to be incorporated. Because a classification method usually does not make strong assumptions about the inputs, this also allows us to model the relationships between multiple ChIP-seq samples without specifying their distributions. In theory, any classification method can be used here but typically an ANN is used, which allows flexible modeling of the inputs. To train an ANN, the pairs of 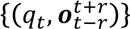 are required as training data. In previous applications of this hybrid model, such as ASR, forced Viterbi alignment can be used to generate *q*_*t*_ because the transcriptions of the speech waveforms are generally available. In AGS, we will have to first hypothesize *q*_*t*_ from observations without using any training labels. Here we will use an HMM-GMM as the boot model to generate the initial values for *q*_*t*_. After that, the hypothesized *q*_*t*_ will be used as targets to train an ANN that will be later combined with an HMM to train a so-called HMM-ANN hybrid model. The hybrid model will then be used to estimate *q*_*t*_ again. In ANN training, the quality of targets is a very important factor to its success. We will discuss two approaches for targets generation from an HMM-GMM in the following.

### 2.2 The Forward-Backward algorithm

Since the *q*_*t*_ have to be estimated, there are intrinsic uncertainties about the estimates and this seems to suggest the use of posterior probabilities 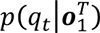 as training labels. The forward-backward algorithm (FB) was previously exploited to provide training labels for ANNs in ASR with success^8^. Here we consider a modified version of it with an additional parameter *w*_*l*_ to adjust the balance between the observation and the transition probabilities. First, let’s give the definition and a recursive formula to calculate the forward and the backward probabilities:

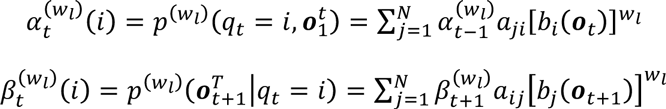

Then, the modified posterior probability can be calculated as

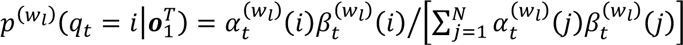

We assign more weight to the observation probabilities than the transition probabilities when *w*_*l*_ > 1, and vice versa when *w*_*l*_ < 1. In this study, *w*_*l*_ is treated as a hyper-parameter and is empirically determined.

### 2.3 The Viterbi algorithm

We can also obtain the 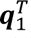 as the most likely state sequence that generates the observations. Similarly, we modify the Viterbi algorithm (VT) that aims to maximize the joint likelihood of 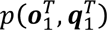. First, assume we already know the maximum of

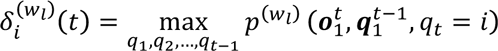

Then, we give a recursive formula 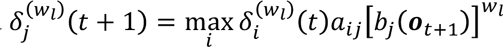 and the initial condition 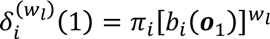 to solve the optimization problem. It shall be noted that the VT is also used in the decoding phase to determine the chromatin state sequence and there is a separate weight parameter *w*_*d*_ to control that, which shares the same definition as *w*_*l*_. Both *w*_*l*_ and *w*_*d*_ are chosen from several values – [0.5, 1.0, 2.0, 3.0, 4.0, 5.0].

### 2.4 Neural network training

In our ANNs, the inputs are the ChIP-seq signals from the neighboring windows centered on each ***o***_*t*_. Each ANN needs to classify the inputs into *N* chromatin states, where *N* is typically larger than 20. We choose to use a softmax output layer to deal with the multi-classification problem and it also serves as a baseline without using any nonlinearity. We then add one layer of hidden units and choose the layer size from a number of values – [250, 500, 1000, 2000, 3000, 4000]. Another layer of hidden units with the same size is added to the ANN to see if there is any improvement. The hidden unit type is chosen to be the rectifier linear function that is found to be fast in convergence and allows a deep ANN to be trained without using pretraining^9^. The ANNs are optimized using stochastic gradient descent (SGD) with momentum and a mini-batch size of 100 samples. The momentum term is initially chosen to be 0.5 and linearly increases to 0.9 after 100 epochs. We initially searched through several values for the learning rate, [1.0, 0.5, 0.1, 0.05, 0.01, 0.005, 0.001] and found 0.01 to be the fastest in convergence without diverging from taking too large steps. The cross-entropy score on a validation set is used to judge stopping criterion. A window of 400 epochs is used to ensure convergence. Dropout is used to prevent overfitting with dropping rates of 0.5 for hidden units and 0.2 for inputs. We choose the radius to be 5 or 10 and the motivation is that 5 consecutive windows correspond to a 1Kb genomic region that is typically the half-size of a functional element such as the promoter and the enhancer. We use *r* = 10 to see if any improvement can be made by including more neighboring signals.

### 2.5 Benchmark dataset

We choose a ChIP-seq dataset from a previous study^5^, which has a mix of histone modifications, transcription factors and open chromatin accessibility. The dataset consists of 31 ChIP-seq samples that were originally used to predict 25 chromatin states at single-base resolution from the human K562 cell line. Here we use the same dataset but average the signals for each 200bp window. We perform training and hyper-parameter tuning on the ENCODE region that is ~1% of the human genome and then make predictions on the whole genome. The ENCODE region contains 149,781 windowed samples out of which 20,000 are used as a validation set and the rest are used as a training set for the ANNs. To evaluate the algorithm’s performance, we use the TSSs with at least two CAGE counts as true positives. A hit is defined as an overlap with at least 1bp between a predicted TSS state and a CAGE-supported TSS, based on which the precision and recall scores are calculated. Two previous state-of-the-art methods, namely ChromHMM and Segway, are compared with our algorithm. We use ChromHMM to train a model on the ENCODE region with its default parameters and make predictions on the whole genome. For Segway, we directly use the result downloaded from its website. We use the Graphical Models Toolkit^10^ for the HMM-GMM and as a framework for the hybrid model, and Pylearn2^11^ for ANN training on a GPU.

## 3 Result

We find the probabilistic mass of *q*_*t*_ generated by the HMM tends to concentrate more on one chromatin state category if we increase the *w*_*l*_. This is not an issue when the VT is used for targets generation, which always assigns 1 for one category and 0 for the rest. We choose *w*_*l*_ = 0.5 and *w*_*l*_ = 3.0 to test how the choice of ANN models affects their abilities to predict *q*_*t*_ from inputs (see Fig. 1). It can be seen that it is much harder to predict the targets when they are more ambiguous (i.e., *w*_*l*_ = 0.5). Using hidden units helps to reduce the loss function but using two hidden layers does not seem to be necessary. In fact, the 1 hidden layer ANNs perform better than the 2-hidden layer ANNs when *w*_*l*_ = 3.0. There is a strong trend that the ANN favors large layer size. The ANNs always perform the best when the layer size is between 2000 and 4000. We therefore conclude that our inputs contain a large number of relatively simple features in comparison with the features used in other application fields of ANNs, such as computer vision and ASR. We also find *r* = 5 to be always better than *r* = 10, i.e., adding farther neighboring windows does not improve the prediction of *q*_*t*_.

**Fig. 1.**
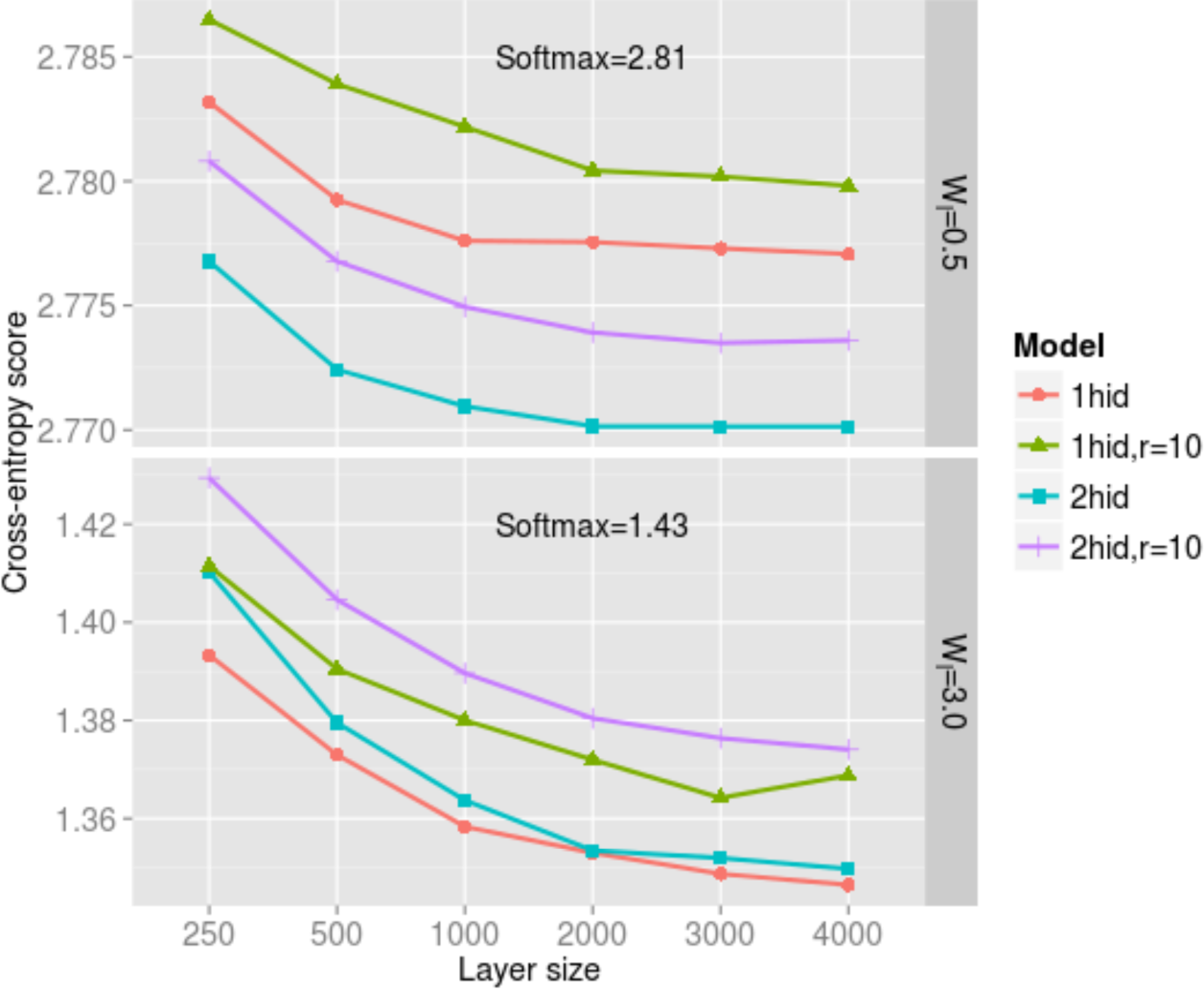
Different ANN models’ Cross-entropy scores for predicting FB generated targets on the validation set.

We then fix the layer size to be 3000, *r* = 5 and use a grid search to find the best combination of *w*_*l*_ and *w*_*d*_ for predicting the TSS state (see Fig. 2). We find the VT generated targets to give robust performance scores against the choice of the weight parameters. While for the FB generated targets, the performance can degenerate when *w*_*l*_ = 0.5, 1.0 or 2.0 depending on the value of *w*_*d*_, no matter a 1-hidden layer or 2 hidden layer ANN is used. We hypothesize that an excess amount of ambiguity in the targets can fail an ANN to capture the regularities among the inputs. Since increased *w*_*l*_ makes the probabilistic mass of *q*_*t*_ to concentrate more on the most promising state, this observation suggests that the VT is a better approach than the FB for targets generation by reducing ambiguities. Table 1 summarizes the performance scores of different models on the whole genome with the weight parameters tuned on the ENCODE region. It can be seen that our HMM-GMM and HMM-ANN models outperform the two existing approaches by large margins. By using the hybrid models, we are able to obtain much higher precisions while maintain similar recalls compared with the HMM-GMM boot model. The hybrid models based on the VT generated targets give the best performance while using a 2-hidden layer ANN does not improve the performance over a 1-hidden layer ANN.

**Fig. 2.**
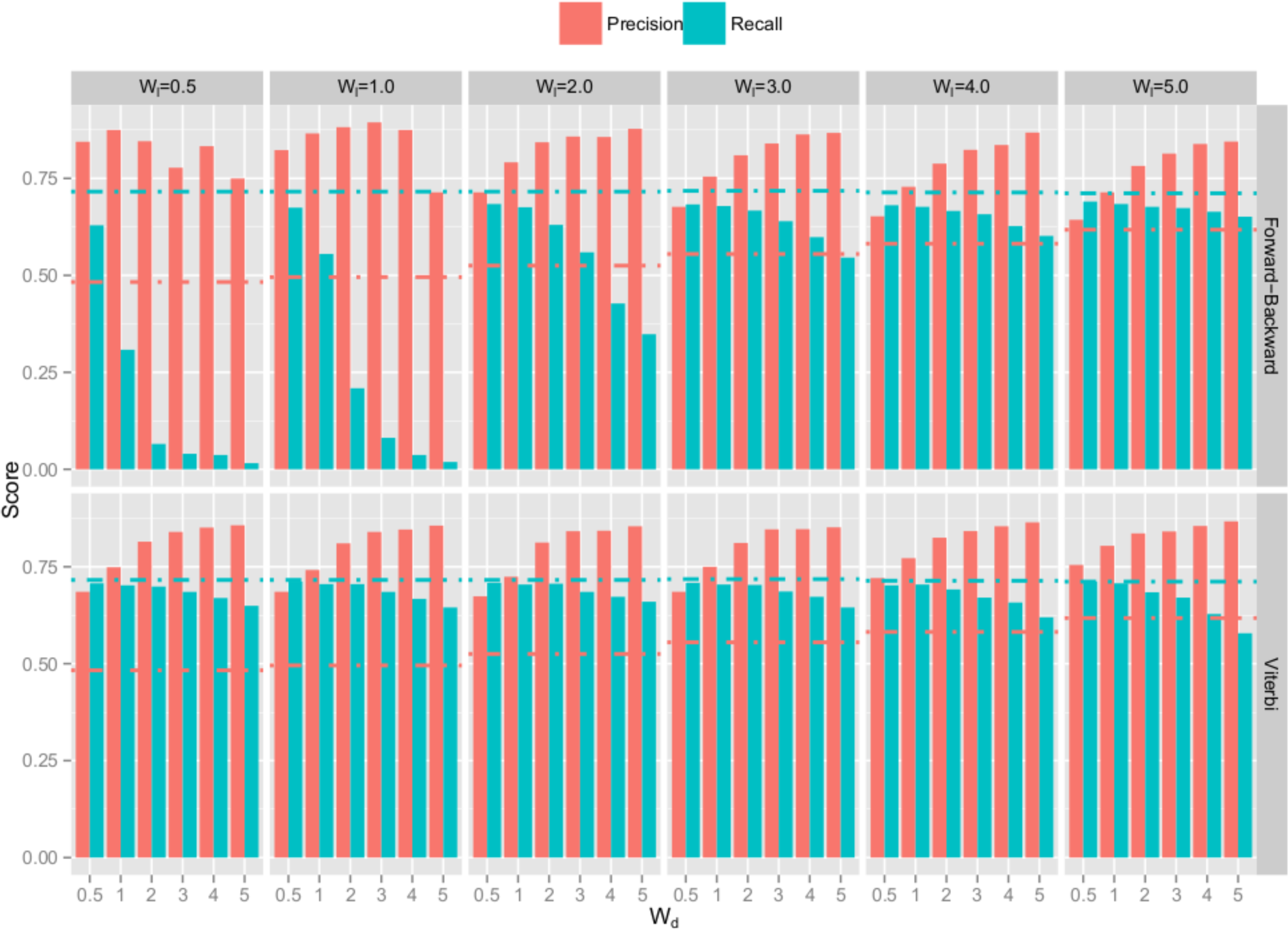
Grid search to determine the target generation and decoding weights on a 1-hidden layer ANN with size of 3000. The dash-dotted lines represent the precision and recall for the HMM-GMM

**Table 1.** Performance scores for different models on predicting the CAGE based TSSs.

| Model | Ref. | Parameters | Precision | Recall | F1 score |
| --- | --- | --- | --- | --- | --- |
| ChromHMM | 4 | Defaults | 0.41 | 0.67 | 0.51 |
| Segway | 5 | Downloaded | 0.32 | 0.73 | 0.44 |
| HMM-GMM | This study | $w_l = 5.0$ | 0.55 | 0.75 | 0.64 |
| HMM-ANN, FB, 1hidx3000 | This study | $w_l = 1.0, w_d = 0.5$ | 0.73 | 0.72 | 0.72 |
| HMM-ANN, VT, 1hidx3000 | **This study** | **$w_l = 3.0, w_d = 3.0$** | **0.74** | **0.73** | **0.73** |
| HMM-ANN, FB, 2hidx3000 | This study | $w_l = 1.0, w_d = 0.5$ | 0.71 | 0.74 | 0.72 |
| HMM-ANN, VT, 2hidx3000 | **This study** | **$w_l = 3.0, w_d = 2.0$** | **0.71** | **0.75** | **0.73** |

